# Specific mutations in SARS-CoV2 RNA dependent RNA polymerase and helicase alter protein structure, dynamics and thus function: Effect on viral RNA replication

**DOI:** 10.1101/2020.04.26.063024

**Authors:** Feroza Begum, Debica Mukherjee, Sandeepan Das, Dluya Thagriki, Prem Prakash Tripathi, Arup Kumar Banerjee, Upasana Ray

**Author notes:** Equal authors (Feroza Begum, Debica Mukherjee, Sandeepan Das, Dluya Thagriki).

## Abstract

The open reading frame (ORF) 1ab of SARS-CoV2 encodes non-structural proteins involved in viral RNA functions like translation and replication including nsp1-4; 3C like proteinase; nsp6-10; RNA dependent RNA polymerase (RdRp); helicase and 3’-5’ exonuclease. Sequence analyses of ORF1ab unravelled emergence of mutations especially in the viral RdRp and helicase at specific positions, both of which are important in mediating viral RNA replication. Since proteins are dynamic in nature and their functions are governed by the molecular motions, we performed normal mode analyses of the SARS-CoV2 wild type and mutant RdRp and helicases to understand the effect of mutations on their structure, conformation, dynamics and thus function. Structural analyses revealed that mutation of RdRp (at position 4715 in the context of the polyprotein/ at position 323 of RdRp) leads to rigidification of structure and that mutation in the helicase (at position 5828 of polyprotein/ position 504) leads to destabilization increasing the flexibility of the protein structure. Such structural modifications and protein dynamics alterations might alter unwinding of complex RNA stem loop structures, the affinity/ avidity of polymerase RNA interactions and in turn the viral RNA replication. The mutation analyses of proteins of the SARS-CoV2 RNA replication complex would help targeting RdRp better for therapeutic intervention.

## 2. Introduction

SARS-CoV2 is a single stranded RNA virus belonging to the family of Coronaviruses [1]. The life cycle of the virus involves receptor attachment and ingress, viral RNA translation, replication, assembly and egress. All these individual steps of the virus life cycle depend on host cell and viral factors.

Viral proteins are of two major categories, the structural proteins and the non-structural proteins. While role of structural proteins is mainly to form the viral structure and encase the viral genome, the non-structural proteins play crucial roles in mediating viral RNA translation and replication [1,2].

In Coronaviruses viral non-structural proteins are encoded by the open reading frame (ORF) 1ab. These include nsp1-4; 3C like proteinase; nsp6-10; RNA dependent RNA polymerase (RdRp); helicase and 3’-5’ exonuclease.

RdRp (also called as nsp12 in SARS-CoV2) is the most important protein helping in viral RNA replication and the helicase helps in unwinding of complex RNA structures so as to help RdRp access the RNA molecule. Proteins are dynamic molecules and their conformations and dynamics might influence their biological functions. Thus, mutations that alter the protein conformations might in turn modulate the dynamics and also the function.

RNA viruses are prone to high frequency of mutations. The rate of mutation itself depends on the fidelity of the RNA polymerase itself and the mutations dictate the virus evolution, immune escape variants and overall variations in the viral genome in the population [3,4]. When polymerase itself is prone to mutation, the fidelity will also get affected and this property might lead to emergence of drug resistant phenotypes. Mutations might also be correlated with the geographical region-specific virulence of a virus variants and carefully targeting the proteins for designing antiviral or vaccine candidates. Along with the structural protein spike, RdRp is another important target of therapeutic intervention.

In this study we have studied the sequence variation of SARS-CoV2 ORF1a polyprotein. We have identified proteins that got mutated the most (RdRp and helicase) and investigated the effect of such mutations on the respective protein structure and function.

## 3. Methods

### 3.1 Sequences

We downloaded 911 sequences of ORF1ab (length: 7096) from the NCBI virus database

### 3.2 Sequence alignments, phytogeny and structure

All the downloaded sequences were aligned by multiple sequence alignment platform of ‘Multiple Alignment using Fast Fourier Transform’ (MAFFT). The alignment file was viewed using Wasabi sequence viewer as well as on JalView and differences in the sequence i.e. mutations recorded.

Phylogenetic tree was generated from the MAFFT alignment file using ‘Archaeopteryx’ platform using neighbour joining method.

CFSSP (Chou and Fasman secondary structure prediction) server was used to predict the secondary structures of SARS-CoV2 RdRp and helicase.

To study the effect of mutation on the tertiary structure of the RdRp and helicase and predicting the impact of mutations on conformation, stability and flexibility, structures of wildtype RdRp (PDB ID: 6M71, Chain A) (Reference sequence: YP_009725307.1) and helicase (ID: 6JYT.1, Chain A) (Reference sequence: YP_009725308.1) from the SWISS model repository [5] (a database of annotated 3 dimensional protein structure models which has been generated by SWISS-MODEL homology-modelling pipeline) were uploaded on DynaMut software (University of Melbourne, Australia) [6]. Change in vibrational entropy between wildtype and mutant proteins; the atomic fluctuations and deformation energies were determined. For atomic fluctuation and deformation energy calculations, calculations were performed over the first 10 non-trivial modes of the molecule.

## 4. Results and Discussion

The study was initiated by acquiring sequence data with respect to the ORF1ab polyprotein of SARS-CoV2. We downloaded 908 available sequences of North American origin. This part of the geographic origin was selected as number of available sequences were maximum from this part of the world. More the number of sequences, more relevant a conclusion would be. All the 908 sequences were subjected to multiple sequence alignment in MAFFT software. The alignment file was downloaded and then uploaded in the Jalview and Wasabi softwares to identify the sites of mutation. We have pointed out only those mutations that we observed to have appeared multiple times i.e. in many isolates. The mutations have been tabulated in Figure 1A.

**Figure 1.**
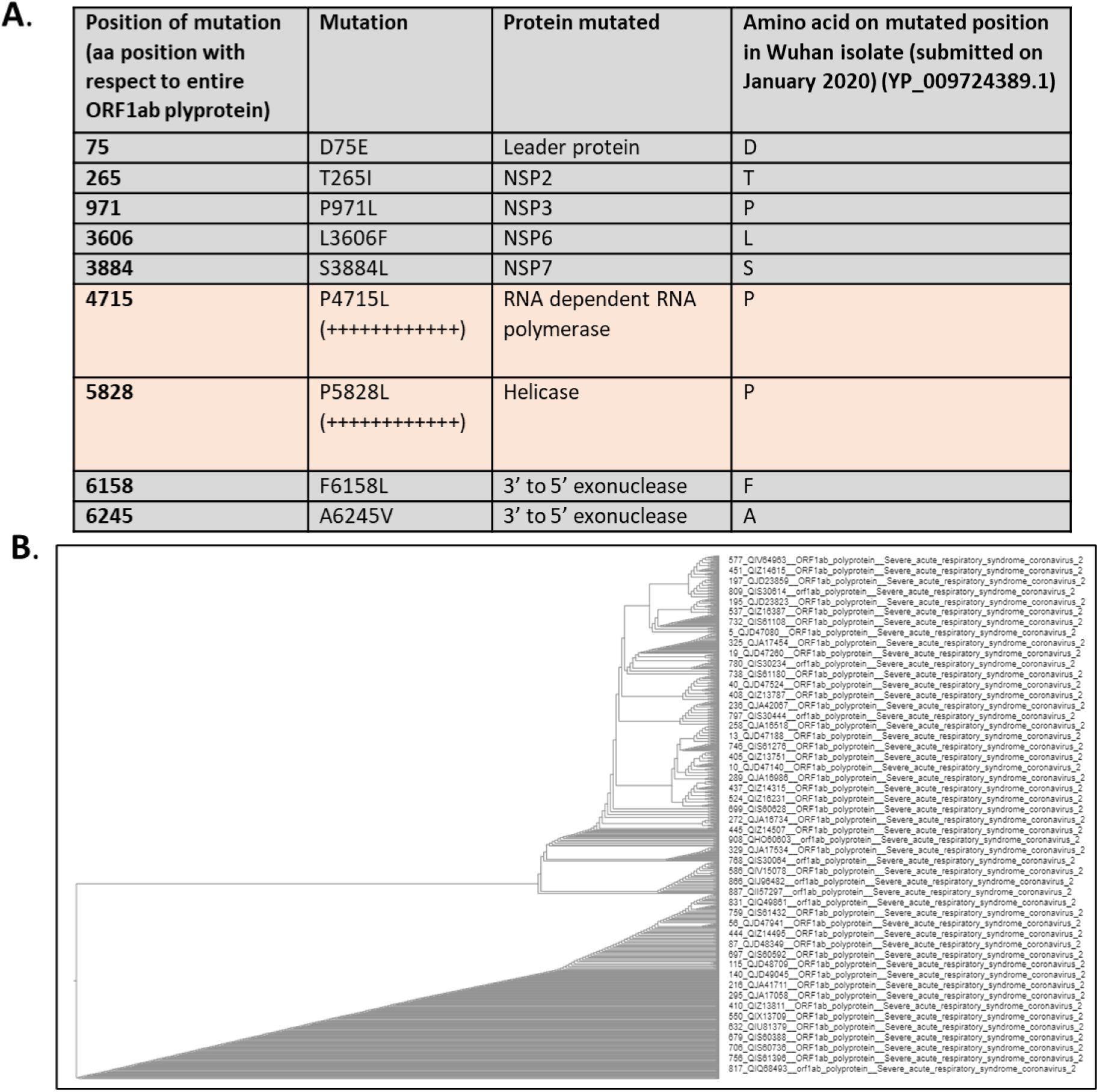
Identification of mutations in the polyprotein of ORFlab and phylogeny. A. Mutations found after MAFFT multiple sequence alignment of ORFlab sequences of 908 isolates B. Phylogenetic tree based on 908 ORFlab sequences

We also compared these mutations with a sequence of Wuhan origin and Indian isolates. However, both Wuhan and Indian isolates didn’t show any of the mutations identified. Phylogeny showed that ORF1ab polyprotein variants formed various clusters showing evolution of this virus in a multipronged form (Fig. 1B).

Out of all the identified prime spots for mutation, the proteins that were seen to be mutated the most are the RdRp and the helicase. Both these proteins are very important for viral RNA replication. RdRp has the polymerase activity and helicase helps in unwinding the tangled RNA for better access of RdRp to the RNA for initiating and carrying out replication. Thus, we further analysed the effect of these mutations on conformation, stability and flexibility of the wild type/ Wuhan type sequence. First, we checked for any possible alterations of the secondary structure of the proteins due to each type of mutation followed by tertiary structure analyses using DynaMut.

In case of RdRp the P to L mutation resulted in loss of two turn structures from positions 323 and 324 in the secondary structure (Figure 2).

**Figure 2.**
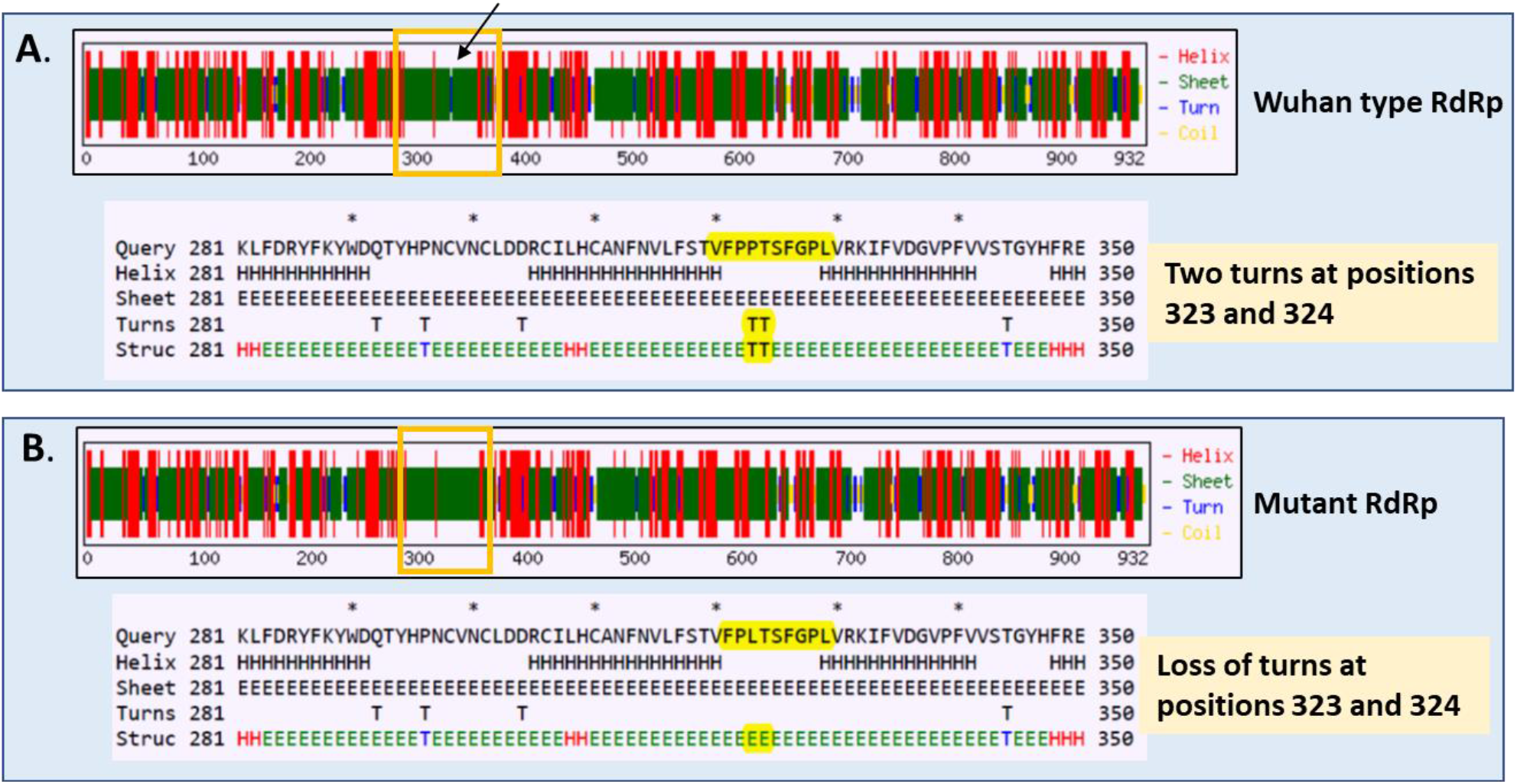
Secondary structure prediction of RdRp. A. Wild-type RdRp. Highlighted part shows position 323 that gets mutated in other isolates B. Mutant RdRp. There is mutation of P to L at position 323. This resulted in loss of two turns at positions 323 and 324

Proline are common in turns and is also called as ‘helix breaker’. On the other hand the amino acid leucine is one of the most stabilizing residues in alpha helix [7]. So, a replacement of proline by leucine is expected to stabilize the structure. The structural analyses revealed that the change in vibrational entropy energy (ΔΔSVib ENCoM) was −4.074 kcal.mol^-1^.K^-1^. There was a decrease in molecular flexibility and ΔΔG was 1.540 kcal/mol (a stabilizing mutation) (Figure 3). Atomic fluctuation and deformation energy calculation studies were performed. Atomic Fluctuation tells the amplitude of the absolute atomic motion. Deformation energy provides a measure for amount of local flexibility in a protein. Calculations were performed over the first 10 non-trivial modes of the molecule (Figure 4). Taken together studies showed that proline to leucine change in RdRp resulted in rigidification of the structure and reduction of molecular flexibility.

**Figure 3.**
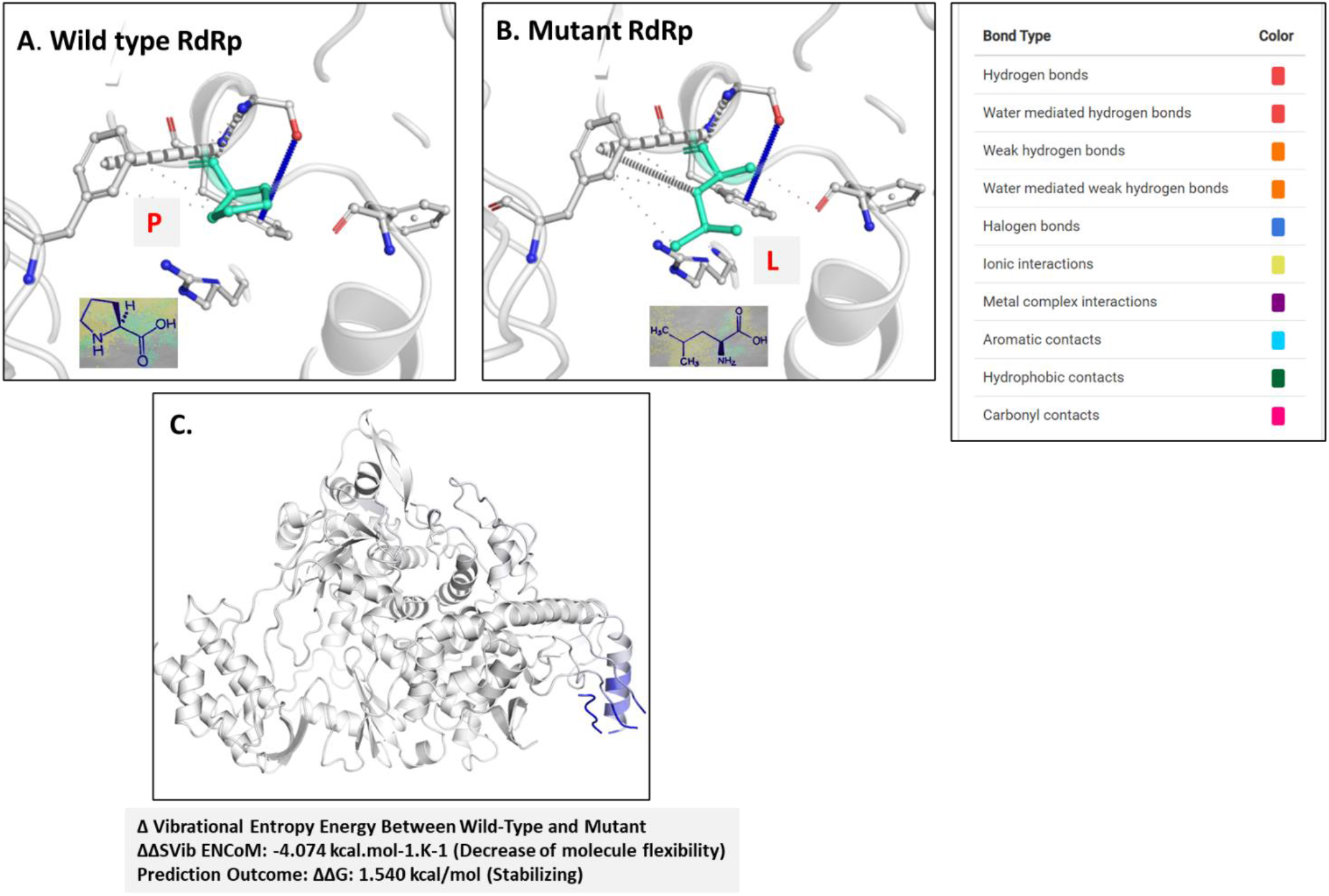
A. Wild-type and B. mutant residues are colored in light-green and are also represented as sticks alongside with the surrounding residues which are involved on any type of interactions. Colour codes are shown on right panel C. Amino acids colored according to the vibrational entropy change upon mutation. Blue colour represents a rigidification of the structure

**Figure 4.**
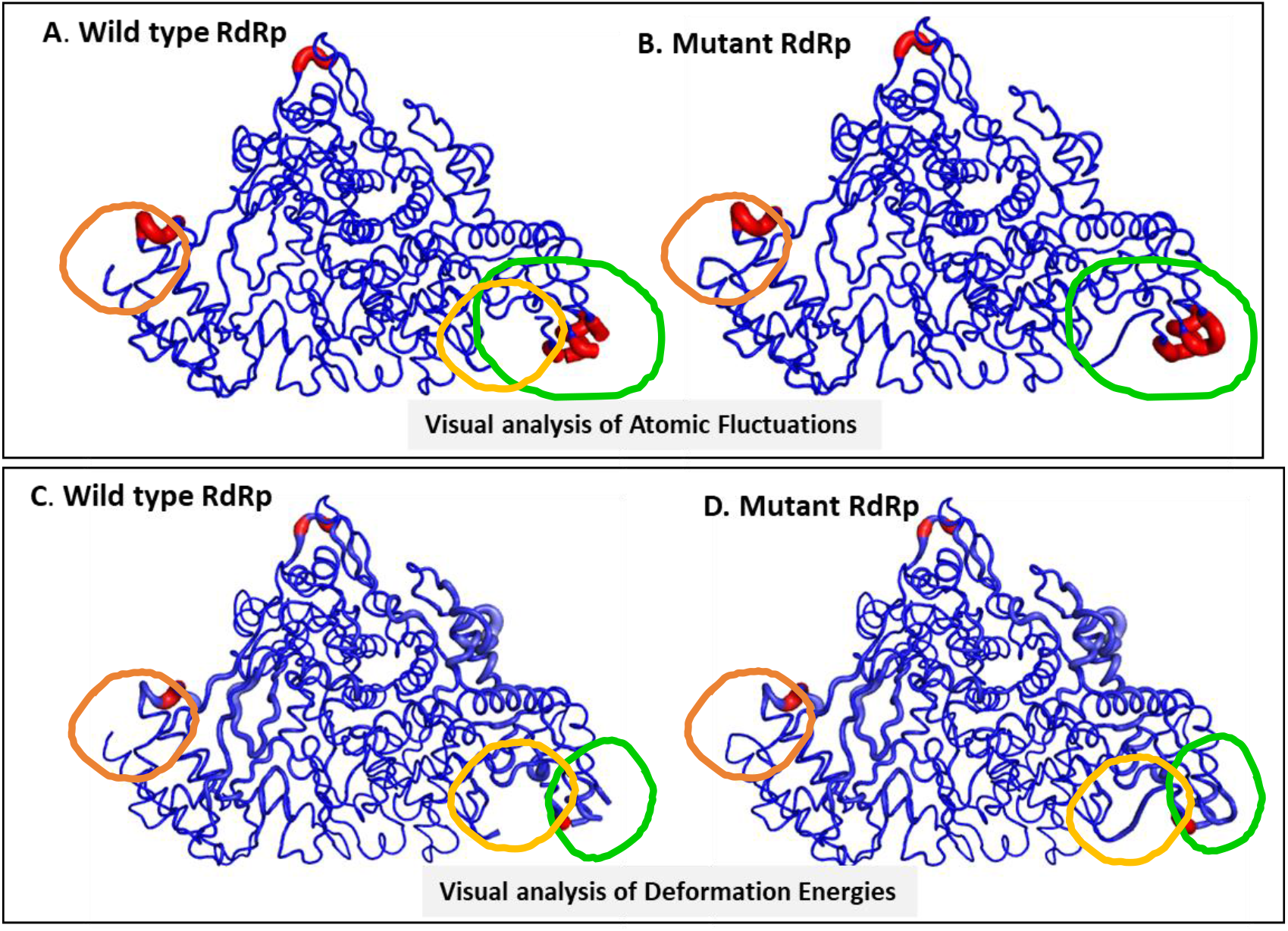
Atomic Fluctuation and Deformation Energy calculations. A. Wild-type and B. mutant RdRp shown after Atomic Fluctuation analyses. Calculations were performed over the first 10 non-trivial modes of the molecule. The magnitude of the fluctuation is represented by thin to thick tube colored blue (low), white (moderate) and red (high). C. Wild-type and D. mutant RdRp after Deformation energy calculations. Calculations performed over the first 10 non-trivial modes of the molecule. The magnitude of the deformation is represented by thin to thick tube colored blue (low), white (moderate) and red (high).

In helicase however, P to L mutation lead to increase in molecular flexibility and the ΔΔG was calculated to be −0.200 kcal/mol and hence destabilizing. Proline is a polar and has uncharged R group whereas Leucine is non polar and has aliphatic R group.

Secondary structure prediction of P to L mutation in helicase revealed significant changes (Figure 5). At positions 503 and 504 there were loss of two turn structures. Three sheets were introduced at positions 502, 503 and 504 of the protein. From positions 500-504, five helices got introduced as compared to the Wuhan type sequence. The DynaMut predictions calculated ΔΔSVib ENCoM to be 0.151 kcal.mol^-^ 1. K^-1^ and an increase in molecular flexibility (Figure 6). Visual analyses of atomic fluctuation and deformation energy calculations are shown in Figure 7.

**Figure 5.**
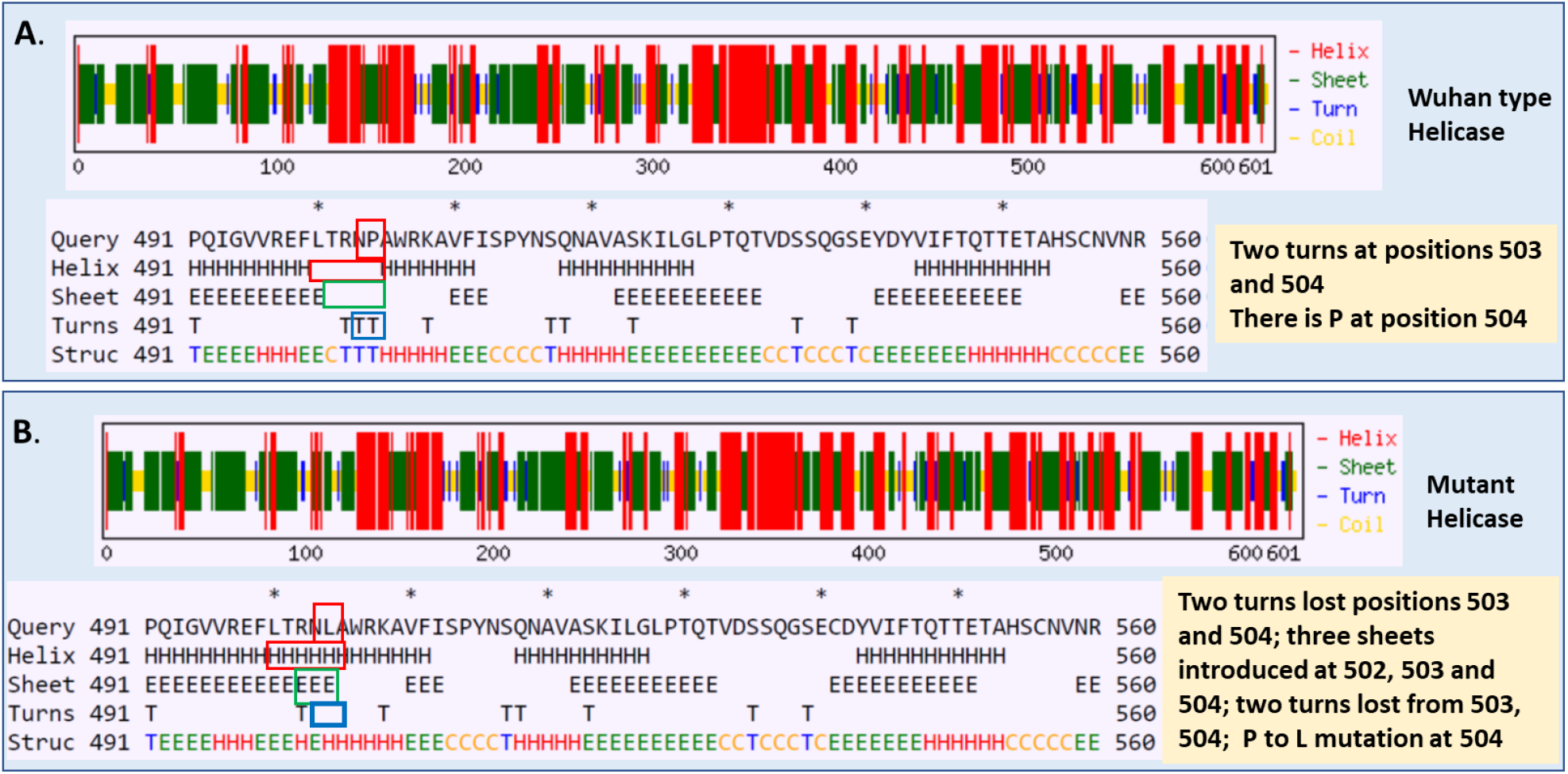
Secondary structure prediction of Helicase. A. W¡ld-tvpe Helicase. B. Mutant RdRp. There is mutation of P to L at position 323. This resulted in loss of two turns at positions 323 and 324

**Figure 6.**
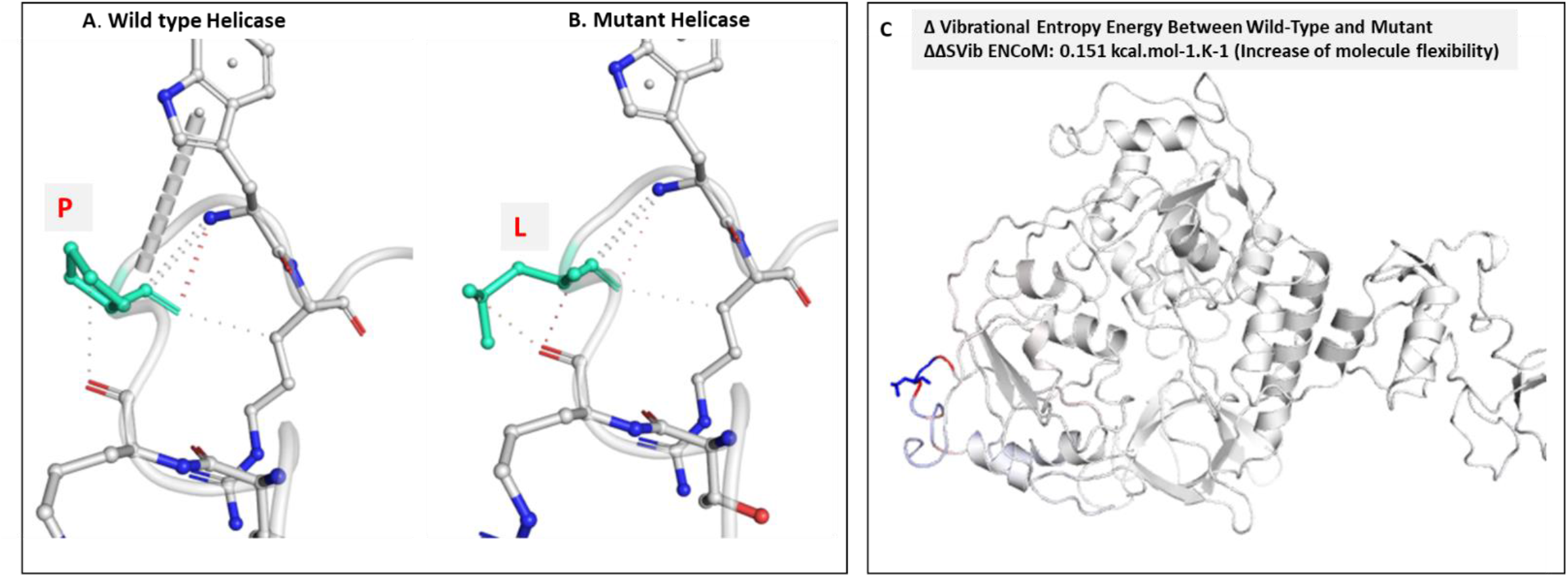
A. Wild-type and B. mutant residues are colored in light-green and are also represented as sticks alongside with the surrounding residues which are involved on any type of interactions. Colour codes are same as shown on right panel in Figure 2 C. Amino acids colored according to the vibrational entropy change upon mutation. Blue colour represents rigidification of the structure and red, a gain in flexibility

**Figure 7.**
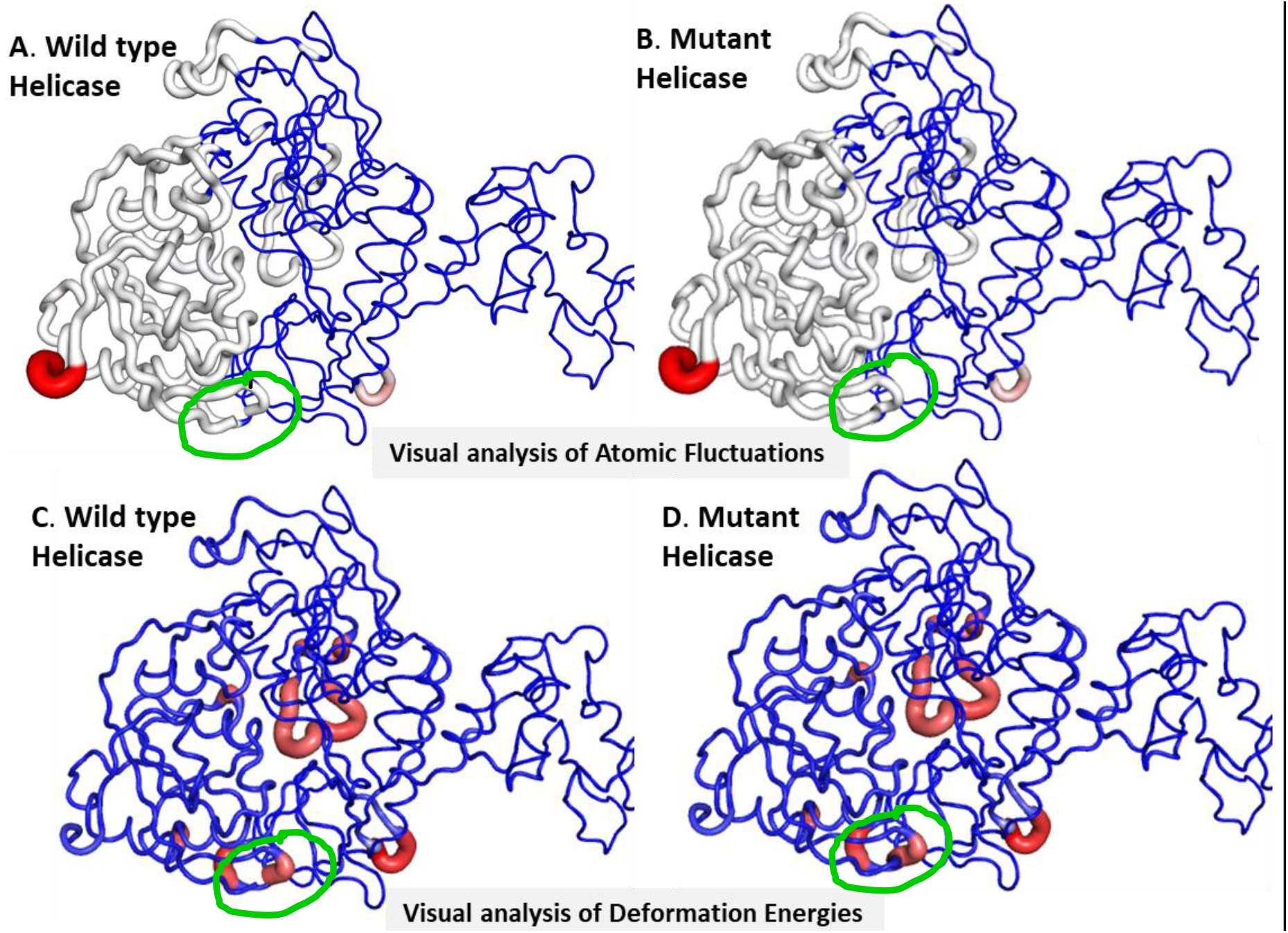
Atomic Fluctuation and Deformation Energy calculations. A. Wild-type and B. mutant helicase shown after Atomic Fluctuation analyses. Calculations were performed over the first 10 non-trivial modes of the molecule. The magnitude of the fluctuation is represented by thin to thick tube colored blue (low), white (moderate) and red (high). C. Wild-type and D. mutant helicase after Deformation energy calculations. Calculations performed over the first 10 non-trivial modes of the molecule. The magnitude of the deformation is represented by thin to thick tube colored blue (low), white (moderate) and red (high).

Although not discussed in detail in the current paper, we also observed two sites in the exonuclease getting mutated in some of the isolates. Mutated exonuclease might be defective of proofreading activity and thus might also influence fidelity [8]. All the three proteins RdRp, helicase and exonuclease are replication determinants and thus mutations in these proteins cam immensely regulate the viral RNA functions.

All these mutations that change the secondary structure and structural flexibility of RdRp is likely to influence the replication of the viral RNA as both these proteins are key players in executing replication of the RNA genome. Fidelity of RdRp is prone to get fine-tuned by mutations in RNA viruses and this property helps in emergence of fidelity variants. High mutation rate might also allow new variants to escape antibodies and establish as antibody escape mutants which might also lead to expansion of tissue tropism of a virus. Both attenuation as well as virulent forms might emerge and influence disease profile.

## Acknowledgements

We thank CSIR, AcSIR and North Bengal Medical College and Hospital for necessary support and input..

